# Liquid-like protein interactions catalyze assembly of endocytic vesicles

**DOI:** 10.1101/860684

**Authors:** Kasey J. Day, Grace Kago, Liping Wang, J Blair Richter, Carl C. Hayden, Eileen M. Lafer, Jeanne C. Stachowiak

## Abstract

During clathrin-mediated endocytosis, dozens of proteins assemble into an interconnected network at the plasma membrane. As early initiators of endocytosis, Eps15 and Fcho1 are responsible for locally concentrating downstream components on the membrane surface. However, they must also permit dynamic rearrangement of proteins within the budding vesicle. How do initiator proteins meet these competing demands? Here we show that Eps15 and Fcho1 rely on weak, liquid-like interactions to efficiently catalyze endocytosis. In reconstitution experiments, these weak interactions promote the assembly of protein droplets with liquid-like properties, including rapid coalescence and dynamic exchange of protein components. To probe the physiological role of liquid-like interactions among initiator proteins, we tuned the strength of initiator protein assembly in real time using light-inducible oligomerization of Eps15. Low light levels drove initiator proteins into liquid-like assemblies, restoring normal rates of endocytosis in mammalian Eps15 knockout cells. In contrast, initiator proteins formed solid-like assemblies upon exposure to higher light levels. Assembly of these structures stalled vesicle budding, likely owing to insufficient molecular rearrangement. These findings suggest that liquid-like assembly of early initiator proteins provides an optimal catalytic platform for endocytosis.

---

Clathrin-mediated endocytosis (CME) is driven by the assembly of a highly interconnected network of adaptor proteins that participate in cargo selection, clathrin recruitment, and membrane bending. A small subset of adaptor proteins is the first to arrive at endocytic sites^1,2^. These initiators include the Fcho proteins (Fcho1/2), Eps15/Eps15R, and intersectin, which associate together in a complex thought to serve as a membrane-associated scaffold for recruitment of downstream adaptors^3^. Specifically, Fcho and Eps15 work together to recruit AP-2, which promotes maturation of the nascent endocytic structure^4–6^. Fcho also binds the adaptors HRB and Dab2^7,8^. Eps15 interacts with numerous other adaptors, including Epsin and AP180/CALM^9,10^. Meanwhile, the minimal initiator complex itself is highly interconnected. Fcho binds Eps15 through interactions between its C-terminal μHD and motifs in Eps15’s intrinsically disordered C-terminus^6,11^. Fcho exists as a homodimer, mediated by its F-BAR domain^12^. Eps15 not only dimerizes via its coiled-coil domain, but can likely also take on an antiparallel tetrameric conformation mediated by interactions between its N-terminal EH domains and C-terminal disordered region^11,13^. Therefore, each Fcho dimer can interact with multiple Eps15 oligomers, suggesting an interconnected initiator protein network. Loss of both Fcho1 and Eps15 results in less productive CME^3,6^.

As initiators of endocytosis, the network formed by Fcho and Eps15 must meet the competing demands of (i) locally concentrating downstream adaptors, while (ii) permitting them to rapidly rearrange as the endocytic structure grows and changes^14,15^. Interestingly, Fcho and Eps15 are thought to be pushed to the edges of the growing pit, where they are ultimately excluded from the budding vesicle^16,17^. This finding illustrates the importance of dynamic rearrangements during the initiation of CME and also suggests that Fcho and Eps15 can be thought of as “catalysts” of endocytosis, because they accelerate the assembly of vesicles but are left behind as the vesicle “product” departs from the membrane.

Here we set out to determine the physical properties that make the initiator proteins effective catalysts. Our results reveal that a network of weak, liquid-like interactions among initiator proteins maximize the productivity of CME. Specifically, using purified proteins in vitro, we find that Fcho1 and Eps15 assemble together into liquid-like protein droplets. Formation of these droplets requires multivalent interactions between Fcho1 and Eps15. In particular, assembly depends strongly on the homo-oligomerization of Eps15 through its coiled-coil domain^18^. Knockout of Eps15 in mammalian epithelial cells results in a substantial increase in the fraction of short-lived, abortive endocytic events, consistent with previous findings^19,20^. We replaced Eps15 in these cells with a chimeric form in which the coiled-coil dimerization domain was replaced by the light-activated oligomerization domain of the protein Cry2. This chimera provided tunable control over the strength of initiator protein interactions. Application of low light to these cells restored wild-type levels of productive endocytosis. In contrast, exposure to stronger light drove solid-like assembly of initiator proteins, leading to stalled, unproductive endocytic structures. Together these results suggest that liquid-like assembly provides a balance between the competing roles of initiator protein catalysts–to facilitate both strong assembly and dynamic rearrangement of growing endocytic structures.

## Results

### Initiator proteins co-assemble on membrane surfaces

The endocytic adaptors Eps15 and Fcho1 are among the earliest proteins to arrive on the membrane during clathrin-coated vesicle formation^1^. To understand how these proteins function together during endocytic initiation, we first examined their assembly on the surfaces of giant unilamellar vesicles (GUVs). Fluorescently labeled full-length proteins at a concentration of 500 nM were introduced to GUVs containing PtdIns(4,5)P_2_. Individually, both Eps15 and Fcho1 decorated these GUVs homogeneously (Figure 1a, Supplementary Figure 1a). Notably, while binding of Eps15 to PtdIns(4,5)P_2_ -containing GUVs was weak^3^, Eps15 bound homogeneously even when recruited more strongly via a histidine tag to GUVs containing NTA-Ni (Figure 1a). In contrast, when both proteins were added simultaneously, they co-partitioned into protein-rich and protein-poor domains on the membrane surface, which displayed strong co-localization between Eps15 and Fcho1 signals (Figure 1b). The protein-rich domains displayed various patterns on the GUVs surfaces, typically consisting of several bright, rounded regions (Figure 1b) or a single bright region (Figure 1c). By comparison, a membrane dye, TR-DHPE, maintained a relatively homogenous distribution, suggesting a lack of long-range heterogeneity in the underlying membrane (Figure 1d). Over 70% of GUVs exposed to both full-length Eps15 and Fcho1 displayed protein-rich domains, while domains were observed on fewer than 10% of GUVs exposed to either protein individually.

**Figure 1.**
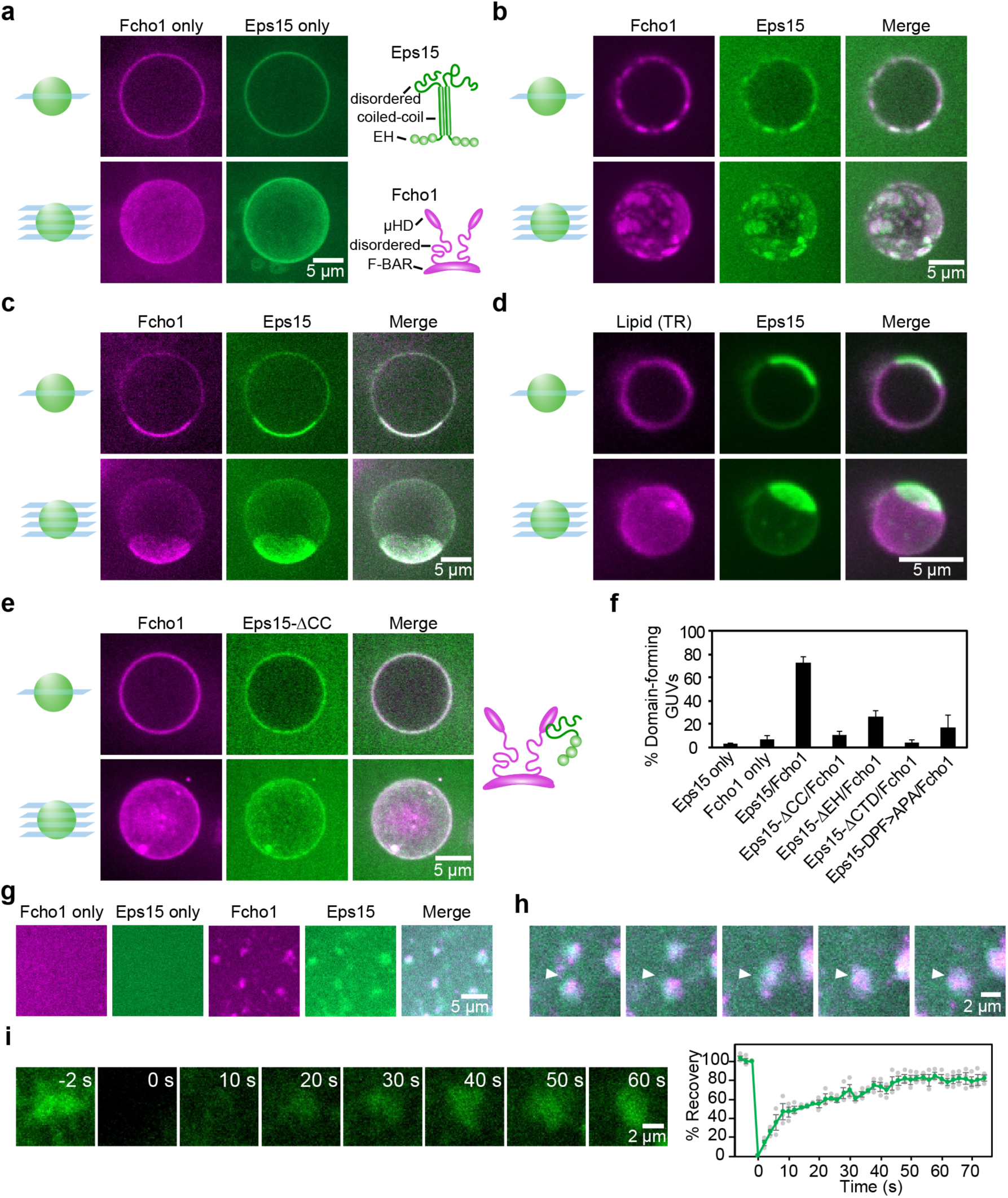
Eps15 and Fcho1 assemble into protein-rich domains on membrane surfaces. (**a-e**) Center slices (upper panels) and corresponding z-projections (lower panels) of representative GUVs incubated with 500 nM of the indicated protein(s). Fcho1 is labeled with Atto-594 and Eps15 variants are labeled with CF488a. GUVs contain 79% DOPC, 15% DOPS, 5% PtdIns(4,5)P_2_, and 1% DPEG10-biotin unless otherwise indicated. Scale bars are 5 μm. (**a**) Full-length Fcho1 alone on GUVs, (left) and full-length Eps15 alone on GUVs containing 97% DOPC, 2% DOGS-NTA-Ni, 1% DPEG10-biotin (center). Cartoons (right) depict domain organization of Fcho1 and Eps15 dimeric forms. (**b**) GUVs incubated with both Fcho1 and Eps15 display several protein-rich domains or (**c**) a single protein-rich domain. (**d**) GUVs labeled with 0.1% Texas Red-DHPE lipid were incubated with 500 nM each of CF488a-labeled Eps15 and unlabeled Fcho1. (**e**) GUVs incubated with Fcho1 and Eps15 lacking the coiled-coil domain (Eps15-ΔCC) do not display protein-rich domains. Cartoon depicts interaction between Fcho1 and an Eps15-ΔCC monomer. (**f**) Frequency of GUVs displaying protein-rich domains for each set of proteins. For each bar, n=4 experiments with at least 50 total GUVs for each condition, error bars show SEM. (**g-i**) Multibilayers containing 73% DOPC, 25% DOPS, and 2% DOGS-NTA-Ni were incubated with 100 nM Atto594-labeled Fcho1, or 100 nM CF488a-labeled Eps15, or both. Scale bar is 5 μm. **(g)** Fcho1 and Eps15 individually decorate multibilayers homogeneously. When combined, Fcho1 and Eps15 form micron-scale protein-rich regions. Scale bar is 5 μm. (**h**) Time course of protein-rich Eps15/Fcho1 domains merging on a multibilayer, 6 s intervals. Scale bar is 3 μm. **(i)** Representative images and plot of fluorescence recovery after bleaching Eps15 (green)/Fcho1 (unlabeled) protein domains on multibilayers. n=3 experiments, individual data points are shown in gray. Error bars show SEM.

How does the assembly of protein-rich and protein-poor domains depend upon interactions between Eps15 and Fcho1? Eps15 contains three N-terminal EH domains, followed by a dimerizing coiled-coil domain and a ∼358 amino acid C-terminal intrinsically disordered region^3,11,16^ (Figure 1a). Interspersed throughout the disordered region are 15 tripeptide Asp-Pro-Phe (DPF) motifs, which bind other clathrin adaptors as well as the EH domains of Eps15^21,22^. When exposed to the combination of Fcho1 and a version of Eps15 lacking the coiled-coil domain (Eps15-ΔCC), protein domains were observed on fewer than 10% of GUVs (Figure 1e,f). Deletion of other portions of Eps15 also impacted the formation of protein-rich domains. Specifically, removal of the three N-terminal EH domains (Eps15-Δ3xEH) resulted in slightly weaker partitioning of the initiators into protein-rich domains (Figure 1f, Supplementary Figure 1b). In contrast, deletion of the C-terminal disordered region (Eps15-ΔCTD) blocked the recruitment of Eps15 by Fcho1 to PtdIns(4,5)P_2_ membranes, preventing domain formation (Supplementary Figure 1c). A binding interaction between Eps15 and the μHD of Fcho1 has been mapped previously to a stretch containing three DPF motifs (amino acids 623-636) within Eps15’s disordered region^6^. To test the importance of these residues for Eps15/Fcho1 assembly on the membrane, we mutated each of the three motifs to Ala-Pro-Ala (Eps15-DPF>APA). Interestingly, this Eps15 mutant was still recruited to the membrane by Fcho1 and drove assembly, albeit more weakly than wild-type Eps15 (Supplementary Figure 1d). The ability of Eps15-DPF>APA, but not Eps15-ΔCTD, to drive assembly into domains with Fcho1 indicates an important role for the disordered region. Taken together, these results indicate that several multivalent interactions are required for assembly of Eps15 and Fcho1 into an extended network on the membrane surface in vitro.

To probe the dynamics of the protein-rich domains, it was helpful to use a flat membrane substrate, so we observed the assembly of Eps15 and Fcho1 on the surfaces of lipid multibilayer stacks adhered to coverslips^23^. As on GUVs, Eps15 and Fcho1 bound uniformly to multibilayers when applied individually (Figure 1g). However, when applied together at a concentration of 100 nM each, the two proteins rapidly condensed into bright micron-scale domains (Figure 1g). When domains came into contact with each other, they readily merged together (Figure 1h, Supplementary Video 1), suggesting fluid-like assemblies. Additionally, the domains recovered rapidly after photobleaching (Figure 1i), indicating rapid molecular exchange. Together these data suggest that assemblies of Eps15 and Fcho1 bear some of the hallmarks of liquid-liquid phase separation, including (i) substantial regions of intrinsic disorder^24,25^, (ii) the ability to form an extended network based on weak multivalent interactions^26,27^, (iii) fluidity, and (iv) rapid molecular exchange with molecules in solution^28^. However, from these data alone it is unclear whether assemblies of Eps15 and Fcho1 display the thermodynamic properties of a two-phase liquid system^29,30^, such as uniform dissolution of the droplet phase above a well-defined critical temperature. To more directly address these questions, we next characterized the protein assemblies in the absence of the membrane.

### Initiator proteins form liquid-like droplets in solution

We first examined a solution of full-length Eps15 at physiological pH and salt (pH 7.5, 150 mM NaCl), adding 3% polyethylene glycol (PEG) to mimic the crowded cellular environment^31^. At protein concentrations of 4 μM or higher, Eps15 alone readily assembled into large, rounded droplets (Figure 2a). Upon contact, these droplets fused together and re-rounded within 1-2 seconds (Figure 2b, Supplementary Video 2), indicating a liquid-like protein assembly. Furthermore, fluorescence recovery after photobleaching (FRAP) of Eps15 droplets revealed rapid exchange of proteins between droplets and the surrounding solution, with a mobile protein fraction of 80±5% (SEM, Figure 2c).

**Figure 2:**
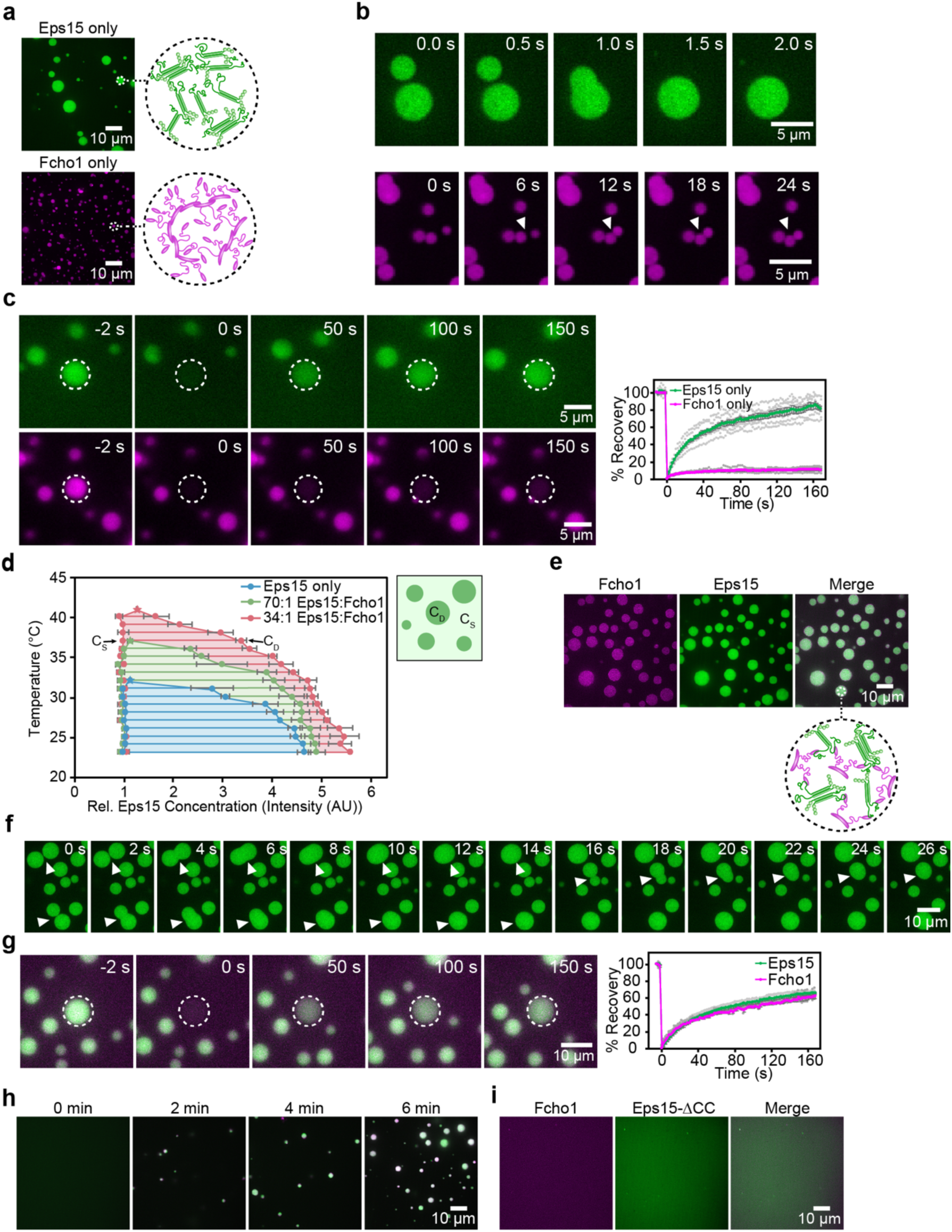
Eps15 and Fcho1 co-assemble into liquid-like protein droplets. Fcho1 is labeled with Atto-594, Eps15 or Eps15-ΔCC is labeled with CF488a. All droplet experiments are at physiological pH and salt with 3% PEG. (**a**) 7 μM Eps15 forms large, rounded droplets, 7 μM Fcho1 clusters into small, irregular aggregates. Insets show cartoon of inferred protein network assembly within droplets. (**b**) Time course of Eps15-only droplets (upper panels) undergoing fusion (arrowheads) and Fcho1-only aggregates (lower panels) approaching each other but failing to fuse. (**c**) Representative images of fluorescence recovery after bleaching an Eps15-only droplet (upper panels) and an Fcho1-only droplet (lower panels). Plot displays fluorescence recovery curves for each. n=6 experiments, individual data points are shown in gray. Error bars show SEM. (**d**) Phase diagram of Eps15/Fcho1 droplets mapped by CF488a-labeled Eps15 fluorescence intensity. Stars denote critical points for each set of Eps15:Fcho1 ratios. C_S_ and C_D_ indicate the concentration of Eps15 in solution and in droplets, respectively. Horizontal tie lines connect C_S_ and C_D_ for a given temperature. Total protein was held constant at 7 μM. n=3 experiments, error bars show SEM. (**e**) When combined at a 1 to 34 ratio, Fcho1 and Eps15 co-localize in protein droplets. Inset shows cartoon of inferred protein network assembly within droplets, Fcho1 in magenta and Eps15 in green. (**f**) Time course of three fusion events (arrowheads) between droplets containing unlabeled Fcho1 and labeled Eps15. (**g**) Representative images and plot of fluorescence recovery after bleaching a Fcho1 and Eps15 droplet. n=4 experiments, individual data points are shown in gray. Error bars show SEM. (**h**) Time course of protein phase separation induced by the addition of Fcho1 to Eps15. (**i**) At a 1 to 34 ratio, Fcho1 and Eps15-ΔCC do not co-assemble into droplets.

To probe the thermodynamic properties of Eps15 droplets, we examined how the protein partitioned between the droplet and solution phases while varying temperature. Here the relative fluorescence intensity of the droplets compared to the surrounding solution provides an estimate of the relative protein concentration in the two phases, enabling us to map a temperature-concentration phase diagram. At each temperature, the relative concentrations of Eps15 in solution (C_S_) and in the droplets (C_D_) were plotted and connected by a tie line^30^ (Figure 2d, blue). As the temperature increased, Eps15 droplets gradually diminished in brightness, indicating a decrease in the concentration of protein inside them, relative to the surrounding solution (Supplementary Figure 2). Eps15 droplets dissolved at a critical temperature of approximately 32°C. Furthermore, droplets re-formed when a heated single-phase solution of Eps15 was cooled to below 32°C (Supplementary Video 3). As this temperature is below 37°C, Eps15 alone appears insufficient to drive protein phase separation at the physiological temperature range of mammalian cells. Therefore, we examined the impact of Fcho1 on the critical temperature of phase separation. In these experiments we mapped the phase diagram of Eps15 in the presence of increasing concentrations of Fcho1, while keeping the total protein concentration constant (Figure 2d). In human cells Eps15 is estimated to be one to two orders of magnitude more abundant than Fcho1^32^. When the ratio of Fcho1 to Eps15 was 1 to 70, the critical temperature shifted upward from 32°C to 37°C. Further increasing the proportion of Fcho1 to 1 to 34, the critical temperature shifted still higher to approximately 41°C, above physiological temperature.

The ability of Fcho1 to stabilize liquid condensates of Eps15, both on membrane surfaces (Figure 1a-c) and in solution (Figure 2d), suggests that Fcho1 molecules assemble more strongly in comparison to Eps15 molecules. Indeed we observed that Fcho1 alone assembled into solid-like polymorphous aggregates when observed under the same conditions for which Eps15 alone formed liquid-like droplets (Figure 2a, Supplementary Video 2). In contrast to the rapid fluid-like merging of Eps15 droplets, aggregates of Fcho1 failed to fuse or re-round upon contact (Figure 2b) and displayed a mobile fraction of only 10±1% in FRAP experiments (Figure 2c). Accordingly, droplets consisting of both Fcho1 and Eps15 in an Fcho1 ratio of 1 to 34 (Figure 2e) displayed intermediate dynamics that were more similar to those of Eps15 droplets than to those of Fcho1 aggregates. Co-assembled droplets fused and re-rounded within several seconds (Figure 2f, Supplementary Video 2) and recovered rapidly after photobleaching, with a mobile molecular fraction of 71±2% for Eps15 and 69±7% for Fcho1 (Figure 2g).

If Fcho1 functions to stabilize condensation of Eps15, then it should be possible to trigger assembly of droplets by adding Fcho1 to a solution of Eps15. In agreement with this prediction, 3 μM Eps15 remained soluble in solution at room temperature. However, addition of Fcho1 to a final ratio of 1 to 25 was sufficient to trigger spontaneous co-assembly of both proteins into droplets within minutes (Figure 2h, Supplementary Video 4). Further, we found that Eps15/Fcho1 droplets relied on the same multivalent interactions that were required for assembly of protein domains on membrane surfaces. Specifically, we combined Fcho1 with each Eps15 domain deletion variant at an Fcho1 ratio of 1 to 34. Eps15-ΔCC failed to drive droplet formation alone or in the presence of Fcho1 (Figure 2i). Similarly, Eps15-ΔCTD blocked formation of droplets almost entirely, while Eps15-Δ3xEH was able to drive the formation of very small droplets only in the presence of Fcho1 (Supplementary Figure 3a,b). Finally, the Eps15-DPF>APA mutant assembled into droplets and recruited Fcho1, but the partitioning of Fcho1 into these droplets was weaker than in droplets formed with wild-type Eps15 (Supplementary Figure 3c). Altogether, these results suggest that a multivalent network between Eps15 and Fcho1 is required for assembly of liquid-like condensates at physiological temperature. Importantly, two-dimensional, membrane-bound condensates were observed near physiological protein concentrations^32^ (Figure 1), suggesting that they are more likely to form in cells compared to three-dimensional, membrane-free droplets, which required protein concentrations well above the physiological range (Figure 2).

### Liquid-like assemblies of initiator proteins drive robust endocytosis

How does the liquid-like assembly of Eps15 and Fcho1 observed in vitro impact the function of these proteins in the cell? We hypothesized that the strength of initiator protein assembly might play a role in determining the dynamics of endocytic events. To test this idea, we constructed a chimeric version of Eps15 that was capable of photo-activated assembly. Specifically, we replaced Eps15’s coiled-coil domain, a critical determinant of phase separation (Figure 1e, 2i) with the PHR domain of Cry2 (Figure 3a), which assembles into dimers and oligomers of increasing stability upon exposure to increasing levels of blue light^33^. Similar chimeras between Cry2 PHR and proteins with intrinsically disordered regions have been shown to drive tunable protein droplet formation in the cytoplasm in living cells^34^.

**Figure 3:**
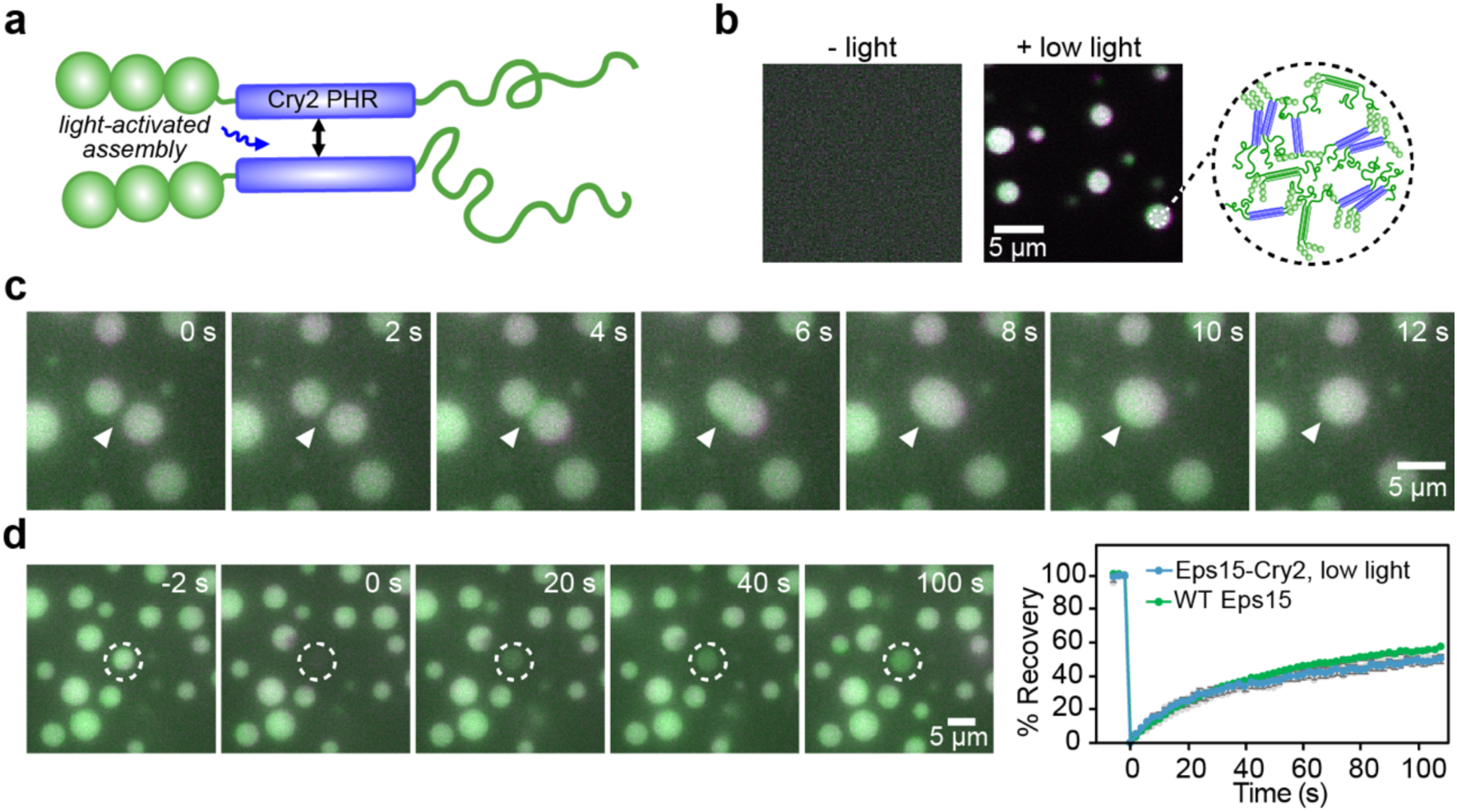
Eps15 can be engineered to assemble into liquid-like protein droplets upon exposure to blue light. (**a**) Diagram of Eps15-Cry2 chimera in which the Eps15 coiled-coil domain is replaced with the Cry2 PHR domain. Blue light exposure drives oligomeric association of Cry2 PHR domains. (**b-d**) Atto594-labeled Eps15-Cry2 and CF488a-labeled Eps15 form droplets after exposure to blue light. (**b**) In the absence of blue light, Eps15 and Eps15-Cry2 are in a single dilute phase (left). Addition of blue light triggers droplet formation (right). Inset shows cartoon of inferred protein network assembly within blue-light induced droplets containing Eps15 and Eps15-Cry2. (**c**) Time course of fusion between droplets containing Eps15-Cry2 and Eps15. (**d**) Recovery of Eps15 fluorescence after bleaching whole Eps15-Cry2/Eps15 droplets is similar to the recovery of Eps15 in WT Eps15/Fcho1 droplets (from Figure 2d). n=3 experiments, individual data points are shown in gray. Error bars show SEM.

We first tested the Eps15-Cry2 chimera in vitro. Our analysis of Eps15 mutant droplet assembly (Figure 2i, Supplementary Figure 3) indicated that a key feature of liquid-like assembly is the requirement for multiple different binding interactions. Therefore, we also incorporated wild-type Eps15 to (i) prevent Cry2 oligomerization from being the dominant interaction driving assembly and (ii) mimic the cellular environment in which multiple different proteins participate in network assembly. At a 1:3 ratio of Eps15-Cry2 to wild-type Eps15, droplets did not form in the absence of blue light. Under these conditions, Eps15-Cry2 appeared to behave similarly to Eps15-ΔCC (Figure 3b, 2i). However, exposure to a low level (10 μW) of blue light induced the formation of droplets, which also incorporated wild-type Eps15 (Figure 3b). These droplets were liquid-like, as they readily merged with one another (Figure 3c, Supplementary Video 5). Further, FRAP of these droplets reflected a high rate of exchange, with t_1/2_=94 s, similar to Eps15 recovery in droplets composed of wild-type Eps15 and Fcho1 (Figure 3d).

We next expressed Eps15-Cry2 in SUM159 human breast cancer-derived epithelial cells. These cells were gene-edited to incorporate a HaloTag on both alleles of the AP-2 σ2 subunit^35^. We used the far-red HaloTag ligand, JF_646_^36^ to mark clathrin-coated structures. We further modified the cells using CRISPR to disrupt expression of both alleles of endogenous Eps15 (Supplementary Figure 4a). We then transiently expressed either wild-type Eps15-mCherry or Eps15-Cry2-mCherry in the resulting Eps15 knockout cells. Cells were imaged using TIRF microscopy, which provides high-contrast visualization of endocytic structures at the plasma membrane^37^.

Both Eps15-mCherry and Eps15-Cry2-mCherry colocalized with labeled AP-2 σ2 in punctate structures (Figure 4a). The lifetime of an endocytic structure on the plasma membrane, measured by the time between the arrival and departure of a labeled protein marker, can vary from tens of seconds to minutes^38^. In general, structures that appear for less than 20 s are considered to be “abortive”, based on previous work demonstrating that they represent transient assemblies that rarely generate productive vesicles^39^. In contrast, “productive” structures, which produce vesicles, typically reside at the plasma membrane from 20 s to a few minutes^38^. Finally, clathrin-coated structures that fail to depart from the plasma membrane within a few minutes are considered “stalled” structures, which are unlikely to represent productive endocytic sites^40^. Each of these three populations appeared in our cells (Figure 4a, b, Supplementary Video 6).

**Figure 4:**
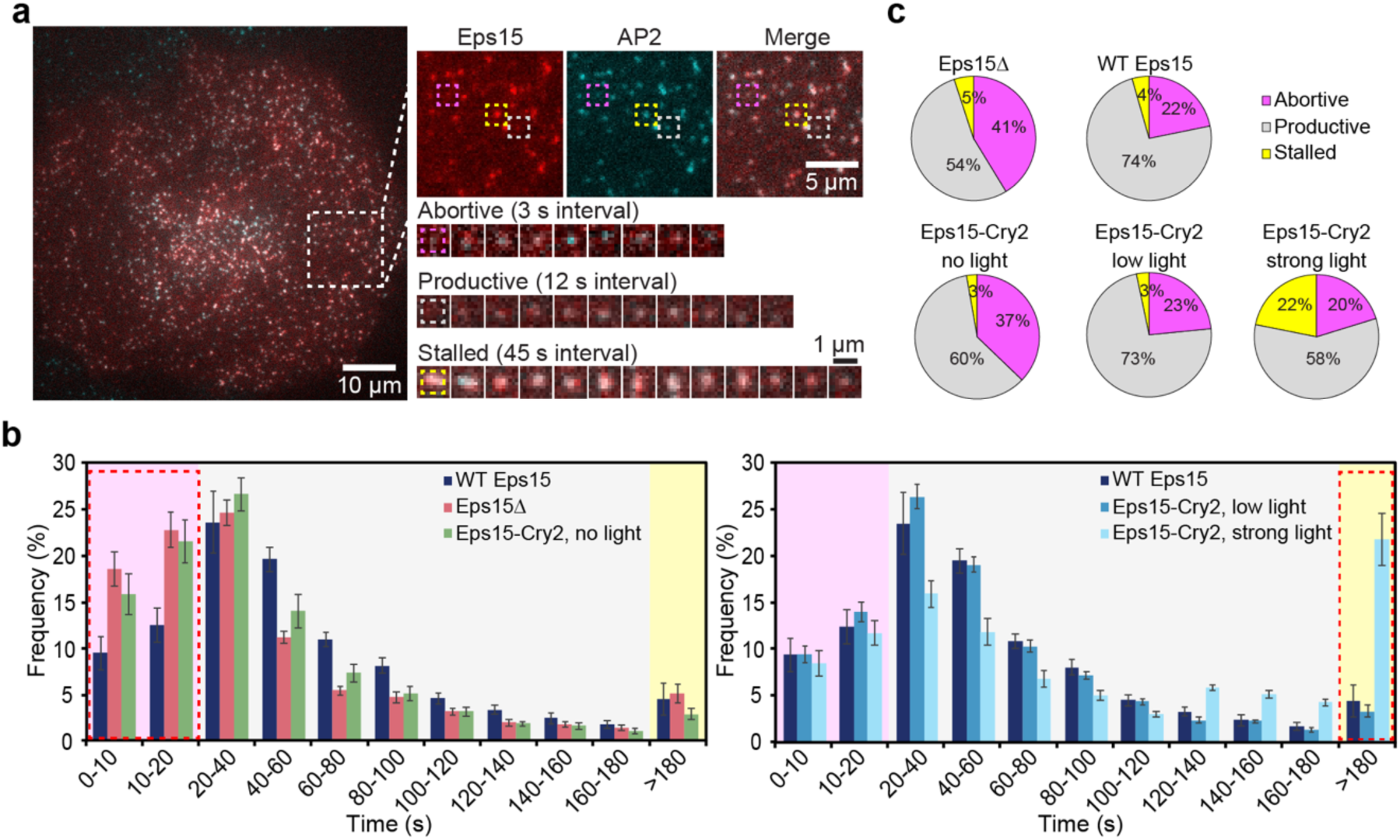
The kinetics of CME can be tuned by light-activated assembly of Eps15. (**a**) Representative image of a cell expressing gene-edited AP-2 σ2-HaloTag conjugated to JF_646_ dye (cyan) and Eps15-mCherry (red). Large inset highlights three representative clathrin-coated structures shown in smaller insets: abortive (magenta), productive (gray), and stalled (yellow) structures lasting 18 s, 96 s, and > 10 min, respectively. (**b**) Histogram of lifetime distributions divided into 11 cohorts of clathrin-coated structures, shaded to indicate lifetimes corresponding to abortive, productive, and stalled structures as in **a**. For each condition, cells lack endogenous Eps15 and express either WT Eps15, no Eps15 (Eps15Δ), or Eps15-Cry2 at no, low, or strong blue light exposure. n=10 or more cells for each condition. Error bars show SEM. (**c**) Lifetime distributions shown in **b** by type of structure.

We first compared cells with endogenous expression of Eps15 to Eps15 knockout cells (Eps15Δ) that were transfected to express Eps15-mCherry. The lifetime distribution of AP-2 σ2-labeled endocytic structures in these two cell populations was nearly identical (Supplementary Figure 4b). Therefore, Eps15Δ cells expressing wild-type Eps15-mCherry were used in subsequent comparisons and will be referred to as WT Eps15 hereafter (Figure 4b, dark blue bars). For all conditions, cells with similar intensity in the mCherry channel were analyzed to maintain comparable expression levels (Supplementary Figure 4c). We then assessed the dynamics of clathrin-coated structures in Eps15Δ cells. Here we observed a higher frequency of short-lived, abortive structures, 41±3% versus 22±3% (Figure 4b, pink bars, 4c). Strikingly, cells expressing Eps15-Cry2 but exposed to no blue light showed a similar sharp increase in abortive structures, to 37±4% (Figure 4b, green bars, 4c). These results highlight the importance of Eps15 assembly for efficient catalysis of CME, building on previous reports that have demonstrated the requirement for multiple domains of Eps15^19,21^.

Next, we analyzed the same Eps15Δ cells expressing Eps15-Cry2, but now with exposure to low (10 μW) levels of blue light. Under these conditions, the frequency of abortive structures was reduced to 23±2%, similar to WT Eps15 cells. Importantly, the low light condition returned the frequency of productive structures to near wild-type levels (Figure 4b,c). These findings demonstrate that light-activated assembly of Eps15 restores the normal kinetics of CME under similar light exposure conditions to those that drove liquid-like assembly of the purified protein in vitro (Figure 3b-d).

### Solid assemblies of initiator proteins stall endocytosis

If a moderate level of blue light can reduce the frequency of abortive structures to wild-type levels, we might expect that exposure to stronger blue light, resulting in stronger assembly of Eps15, would more sharply reduce the number of abortive structures. Indeed, in cells expressing Eps15-Cry2 exposed to strong (50 μW) blue light, the frequency of abortive structures was further reduced to 20±2% (Figure 4c). However, these cells also showed a dramatic increase in the frequency of stalled structures to 22±3%, from 4±2% in WT Eps15 cells, and consequently a reduced frequency of productive structures (Figure 4b, light blue bars, 4c). These results indicate that stronger assembly of the initiator protein network impairs productive maturation of endocytic structures.

To understand the mechanism behind the stall in CME kinetics, we returned to our in vitro analysis of Eps15-Cry2 droplets. We exposed Eps15-Cry2 to strong blue light levels (50 μW) and assessed the physical properties of the resulting droplets. Interestingly, strong blue light exposure drove the assembly of protein droplets that appeared gel-like, as they rarely fused together upon contact and failed to re-round over the course of minutes following contact (Figure 5a, Supplementary Video 5). Further, the droplets displayed weaker recovery after photobleaching in comparison to droplets formed under low levels of blue light (10 μW), with a mobile fraction of 23±2% vs. 52±3% (Figure 5b). Therefore, assembly under strong blue light reduced the dynamic exchange of proteins between the droplets and the surrounding solution. To determine how the strength of the initiator network impacts molecular exchange in live cells, we next analyzed clathrin-coated structures by FRAP (Figure 5c,d). Recovery of wild-type Eps15-mCherry fluorescence occurred with a t_1/2_ of 39 s. In the absence of blue light, Eps15-Cry2 fluorescence recovered substantially more rapidly, with a t_1/2_ of 16 s, suggesting a reduced degree of assembly in comparison to the wild-type protein. Exposure to low levels of blue light (10 μW) returned Eps15-Cry2 fluorescence recovery to wild-type levels, consistent with an increase in assembly of the initiator protein network (t_1/2_=36 s). In contrast, exposure of Eps15-Cry2 to high levels of blue light (50 μW) largely blocked fluorescence recovery, suggesting that endocytic proteins were no longer free to exchange and rearrange within these structures.

**Figure 5:**
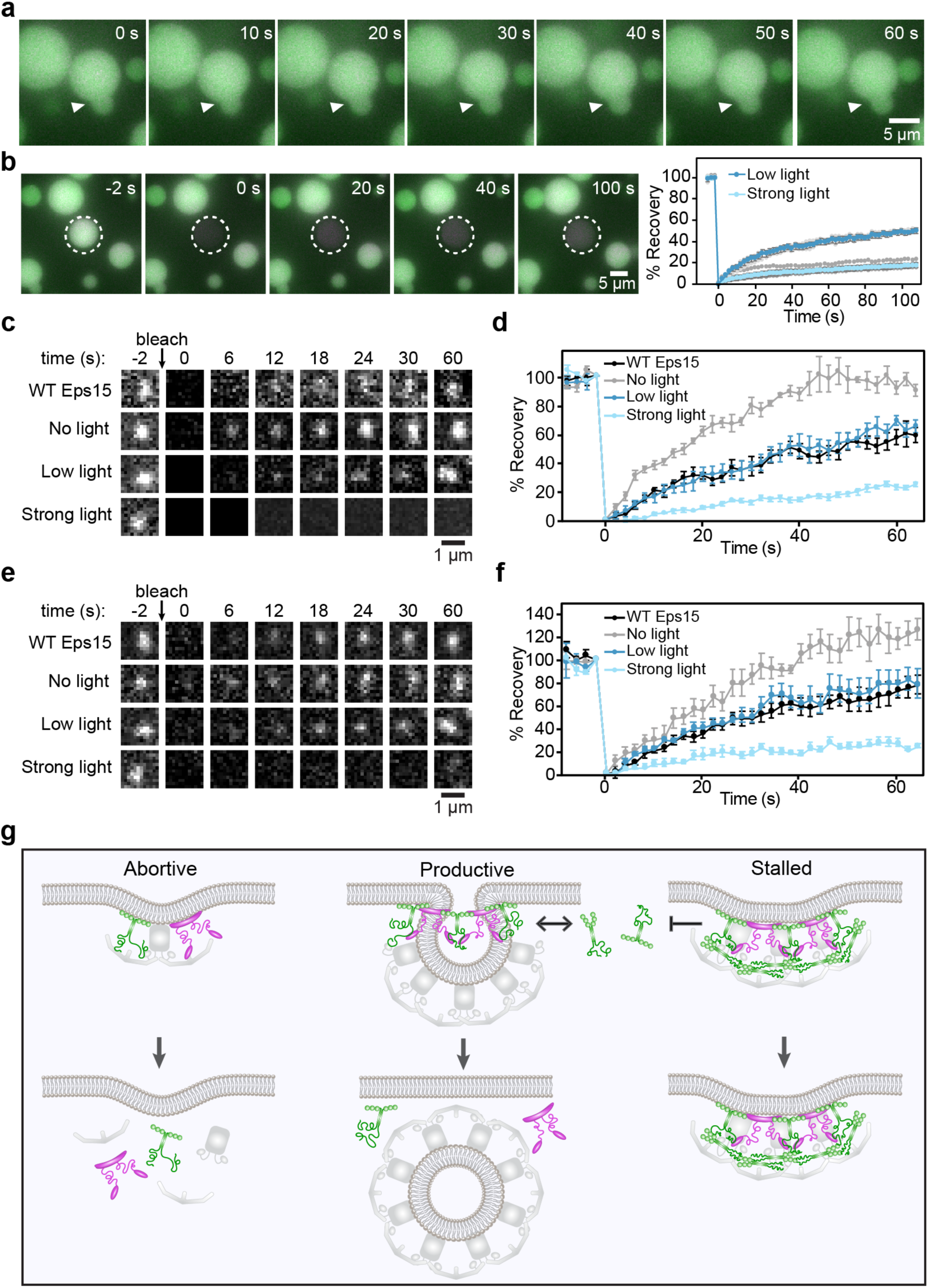
Liquid-like assemblies of initiator proteins optimize the productivity of CME. (**a**) Time course of droplets containing Atto594-labeled Eps15-Cry2 and CF488a-labeled Eps15 failing to merge under strong blue light exposure. (**b**) Recovery of CF488a-labeled Eps15 fluorescence after bleaching an Eps15-Cry2/Eps15 droplet under strong blue light exposure. n= 3 experiments, individual data points are shown in gray. Error bars show SEM. Plot compares recovery to that of the droplet in Figure 4b (Low light). (**c**) Representative images of fluorescence recovery of Eps15-mCherry (top row) or Eps15-Cry2-mCherry in clathrin-coated structures under no, low, or strong blue light exposure. (**d**) Average fluorescence recovery of Eps15 or Eps15-Cry2 for each condition shown in **c**. n=6, error bars are SEM. Plot with individual data points is in Supplementary Figure 4d. (**e**) Representative images of fluorescence recovery of AP-2-HaloTag:JF646 in clathrin-coated structures in cells expressing Eps15-mCherry (top row) or Eps15-Cry2-mCherry under no, low, or strong blue light exposure. (**f**) Average fluorescence recovery of AP-2-HaloTag:JF646 for each condition shown in **e**. n=6, error bars are SEM. Plot with individual data points is in Supplementary Figure 4e. (**g**) Diagram of states of initiator protein assembly in clathrin-coated structures. When Eps15 (green) and Fcho1 (magenta) exist in an unassembled, or dilute phase, abortive structures are favored. In productive structures, Eps15 and Fcho1 assemble into a liquid protein phase capable of exchange with molecules in solution. Further assembly of Eps15 and Fcho1 into a gel or solid phase limits molecular exchange and promotes stalled endocytic structures.

Notably, fluorescence recovery of AP-2, a key cargo adaptor downstream of Eps15 and Fcho^1,6^, showed a similar dependence on blue light exposure in cells expressing Eps15-Cry2 (Figure 5e,f). Specifically, AP-2 recovered to near wild-type levels (t_1/2_=28 s vs. 26 s) in cells exposed to low levels of blue light (10 μW). In contrast, the rate of recovery increased in the absence of blue light (t_1/2_=18 s) and substantially decreased upon exposure to stronger blue light (50 μW). Therefore, the strength of the initiator network controls the freedom of AP-2 to exchange within growing endocytic structures. These results suggest that the interaction strength of the initiator protein network is finely tuned for efficient catalysis of CME. Too little assembly of the initiator proteins produces a structure whose components turn over too rapidly to drive productive endocytic events. In contrast, if the initiator network is too strong, downstream adaptors are unable to rearrange, stalling the growth of endocytic structures. Liquid-like interactions among initiator proteins appear to represent an optimum within this balance.

## Discussion

Despite a detailed biochemical picture of the clathrin adaptor network, the physical mechanisms that drive productive endocytosis have long remained elusive^41,42^. The adaptor network is built from a highly interconnected web of molecular interactions, yet it must undergo extensive remodeling as the endocytic structure matures^15,43^. Here we demonstrate that liquid-like assembly of CME initiator proteins balances these competing constraints, optimizing the productivity of vesicle assembly. Specifically, we have shown that the clathrin adaptor proteins Eps15 and Fcho1 come together in multivalent, liquid-like assemblies, both on membrane substrates (Figure 1) and in 3D droplets (Figure 2). Phase diagrams of these assemblies at varying stoichiometries revealed that both proteins are required in order to generate liquid-like droplets at physiological temperatures (Figure 2d). These results agree with the apparent requirement for both Fcho1 and Eps15 to drive efficient endocytosis^3,6^. Notably, recent work by Kozak and Kaksonen^44^ showed that the *Saccharomyces cerevisiae* homolog of Eps15, Ede1, can also form phase separated condensates in yeast cells, in agreement with our results.

Importantly, binding between Eps15 and Fcho1 is mediated by weak interactions involving short peptide motifs^6^ and intrinsically disordered regions. Similar weak, multivalent interactions are prevalent among CME adaptors^45^. It has been proposed that such interactions facilitate enhanced recruitment of downstream adaptors and promote stability of the adaptor network through avidity^43,46,47^. To probe the requirement for multivalent interactions in cells, we replaced the coiled-coil domain of Eps15 with the Cry2 PHR domain, which provided light-activated control over assembly of the initiator network (Figure 3). In live cells, deletion of Eps15 drove an increase in abortive endocytic events. However, light-induced, liquid-like assembly of the Eps15-Cry2 chimera was able to fully rescue this phenotype. In contrast, solid-like assembly of the chimera, driven by exposure to stronger light, substantially increased the frequency of stalled endocytic events (Figure 4), owing to a loss of molecular exchange within the network (Figure 5c-f).

The initiator protein network has been considered a catalyst of endocytosis because it brings together the key components of the vesicle but is ultimately left behind on the plasma membrane as the endocytic vesicle departs^16,17^. Building on this analogy, all catalysts have two essential functions: (i) to bring together reactants and (ii) to release products. In the context of endocytosis, the coated vesicle, a large supramolecular complex, can be thought of as the product of an assembly reaction, which is catalyzed by the initiator proteins. Interestingly, when the product of a reaction is large, diffusion-limited release from the catalyst often becomes the rate-limiting step^48^. Notably, we observed that productive release of the mature vesicle requires that the initiators are in a liquid-like phase. These results suggest that an essential purpose of the liquid-like assembly of Eps15 and Fcho1 may be to permit diffusion, resulting in efficient release of the mature coated vesicle.

Many pathways of trafficking vesicle biogenesis, including the assembly of COPII vesicles at ER exit sites and clathrin-independent endocytosis, depend upon interwoven protein networks^49–51^. Therefore, liquid-like catalytic complexes, which are capable of balancing the competing demands of multivalent assembly and flexible remodeling, could help to explain the efficient initiation and release of vesicles in trafficking routes throughout the cell.

## Online Methods

### Reagents

Tris-HCl, IPTG, NaCl, β-mercaptoethanol, and Triton X-100 were purchased from Thermo Fisher Scientific. EDTA, EGTA, TCEP, PMSF, protease inhibitor tablets, poly-l-lysine (PLL), Atto 594 NHS-ester and CF488a NHS-ester were purchased from Sigma-Aldrich. Human rhinovirus-3C (HRV-3C) protease, neutravidin, and Texas Red–DHPE were purchased from Thermo Fisher Scientific. Amine reactive PEG (mPEG-Succinimidyl Valerate MW 5000) and PEG-biotin (Biotin-PEG SVA, MW 5000) were purchased from Laysan Bio, Inc. Dipalmitoyl-decaethylene glycol-biotin (DP-EG10-biotin) was provided by Darryl Sasaki. All other lipids were purchased from Avanti Polar Lipids.

### Plasmids

The sequence encoding *M. musculus* Fcho1 was subcloned from Addgene plasmid #27690, a gift from Christien Merrifield^2^, into the pGEX-6P-1 expression vector (GE Healthcare) at EcoRI and NotI restriction sites. pET28a-6xHis-Eps15 (FL), encoding *H. sapiens* Eps15, was a gift from Tom Kirchhausen^11^. Eps15 variant plasmids were derived from pET28a-6xHis-Eps15 (FL). pET28a-6xHis-Eps15-ΔCC was generated by introducing SalI restriction sites to excise residues 315-480 of Eps15. pET28a-6xHis-Eps15-Δ3xEH was generated by PCR amplification to excise residues 15-314 using oligos 5’ ^P^AGTTTACAAAAGAACATCATAGGATCAAGTCC 3’ and 5’ ACTTGATAACTGTGTCAGAGAGAGCTG 3’, followed by ligation. pET28a-6xHis-Eps15-ΔCTD was generated by introducing a NotI restriction site to excise residues 538-896 of Eps15. pET28a-6xHis-Eps15-DPF>APA was generated by site-directed mutagenesis using oligo 5’ GGATTTTTTCCAGTCTGCGCCTGCGGTTGGCAGTGCTCCTGCCAAGGATGCGCCTGCGGGAAAAATCGATCCA 3’ and its reverse complement. To generate both pET28a-6xHis-Eps15-ΔCC::Cry2.PHR and Eps15-ΔCC::Cry2.PHR-pmCherry, the Cry2 PHR domain was PCR amplified from pCRY2PHR-mCherryN1 (Addgene plasmid # 26866, a gift from Chandra Tucker^52^) using oligos 5’ TAGGATCAAGTCCTGTTGCAGCCACCATGAAGATGGACAAAAAGAC 3’ and 5’ ATCAGTTTCATTTGCATTGAGGCTGCTGCTCCGATCAT 3’. This fragment was inserted by Gibson assembly (New England Biolabs) into pET28a-6xHis-Eps15 or Eps15-pmCherryN1 (Addgene plasmid #27696, a gift from Christien Merrifield^2^), which were PCR amplified to exclude Eps15 residues 328-490 using oligos 5’ TCATGATCGGAGCAGCAGCCTCAATGCAAATGAAACTGATGGAAATGAAAGATTTGGAAAATCATAATAG 3’ and 5’ TTGTCCATCTTCATGGTGGCTGCAACAGGACTTGATCCTATGAT 3’.

### Protein Purification

Full-length Eps15, Eps15-ΔCC::Cry2, Eps15-Δ3xEH, Eps15-ΔCC, Eps15ΔCTD, and Eps15-DPF>APA were expressed as N-terminal 6xHistidine-tagged constructs in BL21 (DE3) *E. coli* cells; Fcho1 was expressed as an N-terminal GST-fusion construct in BL21 Star (DE3) pLysS *E. coli* cells. Cells were grown in 2xYT medium for 3-4 hr at 30°C to 0.6-0.9 OD_600_, then cooled for 1 hr before protein expression was induced with 1 mM IPTG at 12°C for 20-30 hr. Cells were harvested, and bacteria were lysed in lysis buffer using homogenization and probe sonication. Eps15-ΔCC::Cry2 was expressed in the dark, protected from direct light, and handled under darkroom red lights during all purification steps.

For Eps15 variants, lysis buffer was 50 mM Tris-HCl, pH 8.0, 300 mM NaCl, 5 mM imidazole, 10 mM β-mercaptoethanol or 5 mM TCEP, 1 mM PMSF, 0.2% Triton X-100, and 1x Roche or Pierce complete EDTA-free protease inhibitor cocktail tablet per 50 ml buffer. Proteins were incubated with Ni-NTA Agarose (Qiagen #30230) resin, followed by extensive washing with 10x column volumes, then eluted from the resin in 50 mM Tris-HCl, pH 8.0, 300 mM NaCl, 200 mM imidazole, 10 mM β-mercaptoethanol or 5 mM TCEP, 1 mM PMSF, and 1x Roche or Pierce complete EDTA-free protease inhibitor cocktail tablet. Full-length Eps15, Eps15-ΔCC::Cry2, Eps15-Δ3xEH, and Eps15-DPF>APA were further purified by gel filtration chromatography using a Superose 6 column equilibrated with 20 mM Tris-HCl, pH 8.0, 150 mM NaCl, 1 mM EDTA, and 5 mM DTT. Purified proteins were concentrated using Amicon Ultra-15 Ultracell-30K centrifugal filter units (MilliporeSigma), then centrifuged at 100k RPM at 4°C for 10 min using a Beckman TLA-120.2 rotor to remove aggregates, and stored either in small aliquots or as liquid nitrogen pellets at −80°C.

For Fcho1, lysis buffer was 100 mM sodium phosphate, pH 8.0, 5 mM EDTA, 5 mM TCEP, 10% glycerol, 1 mM PMSF, 1% Triton X-100, and 1x Roche or Pierce complete EDTA-free protease inhibitor cocktail tablet per 50 ml buffer. Fcho1 was incubated with glutathione sepharose 4B (GE Healthcare), followed by extensive washing with 10x column volumes, then eluted with 15 mM reduced glutathione (Sigma-Aldrich, #G4251) in 100 mM sodium phosphate, pH 8.0, 5 mM EDTA, 5 mM TCEP, 10% glycerol, 1 mM PMSF, and 1x Roche complete EDTA free cocktail tablet per 50ml buffer. Eluted protein was desalted using Zeba Spin Desalting Columns (Thermo Fisher Scientific) into 20 mM Tris-HCl, pH 8.0, 450 mM NaCl, 50 mM KCl, 5 mM EDTA, 5 mM TCEP, and 10% glycerol. The GST tag was cleaved with GST-tagged HRV-3C protease (Thermo Fisher Scientific) overnight at 4°C with rocking. HRV-3C and GST tag were removed by passage through a second GST resin column. Fcho1 was concentrated and centrifuged to remove aggregates as described above and stored as liquid nitrogen pellets at −80°C. All proteins were buffer exchanged into 20 mM Tris-HCl, 150 mM NaCl, 5 mM TCEP 1 mM EDTA, 1 mM EGTA, pH 7.5 before use, except when used on membranes containing DOGS-NTA-Ni, in which case EDTA and EGTA were omitted.

### Protein Labeling

Proteins were labeled using amine-reactive, NHS ester dyes (Atto-Tec) in phosphate-buffered saline containing 10 mM sodium bicarbonate, pH 8.3. The concentration of dye was adjusted experimentally to obtain a labeling ratio of 0.5–1 dye molecules per protein, typically using 2-fold molar excess of dye. Reactions were performed for 15 min on ice, then labeled protein was buffer-exchanged into 20 mM Tris-HCl, 150 mM NaCl, 5 mM TCEP, pH 7.5 and separated from unconjugated dye using Princeton CentriSpin-20 size exclusion spin columns (Princeton Separations). Labeled Fcho1 was centrifuged at 100k RPM using a Thermo Scientific S120-AT3 rotor at 4°C for 10 min to remove aggregates before each use. For all experiments involving labeled protein, a mix of 90% unlabeled/10% labeled protein was used.

### Protein Droplets

All droplets were formed in a buffer of 20 mM Tris-HCl, 150 mM NaCl, 5 mM TCEP 1 mM EDTA, 1 mM EGTA, 3% w/v PEG8000, pH 7.5. For droplet experiments with Fcho1, Eps15, Eps15-ΔCC, Eps15-Δ3xEH, Eps15-ΔCTD, and Eps15-DPF>APA, a total of 7 μM protein was used. Fcho1 and Eps15 variants were combined at 0.1 μM and 6.9 μM, respectively, to achieve an Fcho1 ratio of 1 in 70, or 0.2 μM and 6.8 μM, respectively, to achieve an Fcho1 ratio of 1 in 34. To induce droplet formation in an Eps15 solution by addition of Fcho1, 0.12 μM Fcho1 was added to 3 μM Eps15, yielding a ratio of 1 in 26. Droplets containing Eps15-Cry2 were formed by combining 1 μM Eps15-Cry2 and 3 μM Eps15, then applying 0.5 s of low (10 μW) blue light pulses for 1-3 minutes. For imaging droplets, supported lipid bilayers (SLBs) were used to passivate coverslips before adding droplets solution: small unilamellar vesicles (SUVs) were prepared by drying 100% POPC lipid films in clean glass tubes under vacuum for 2–3 hr. The lipid film was hydrated in 20 mM Tris-HCl, 150 mM NaCl buffer to make a 2 mM liposome solution, which was sonicated with a horn sonicator for 3x 4 min with 2 min cooling intervals while on ice. Wells were prepared by placing 0.8 mm thick silicone gaskets (Grace Bio-Labs) onto ultra-clean coverslips. A 2mM solution of SUVs was added to the coverslip and incubated at room temperature for 15 min to form the SLB. The SLB was washed 6-8x with 20 mM Tris-HCl, 150 mM NaCl, 5 mM TCEP buffer before adding the protein droplet solution.

### GUV Preparation

For Eps15-only experiments, the GUV lipid mixture was 98 mol% DOPC, 2 mol% DOGS-NTA-Ni, and 1 mol% DP-EG10 biotin (for tethering to coverslips). For all other GUVs, the lipid mixture was or 79 mol% DOPC, 15 mol% DOPS, 1 mol% DP-EG10 biotin, and 5mol% PtdIns(4,5)P_2_. Labeled GUVs also incorporated 0.1 mol% Texas Red DHPE. Lipid mixtures were spread in a film on an indium tin oxide-coated glass slide and dried under vacuum overnight. Electroformation was performed at 55°C in 350 milliosmolar glucose solution. The osmolality of the GUV solution and experimental buffer was measured using a vapor pressure osmometer (Wescor). GUVs were tethered to glass coverslips by the following process: wells were made by placing 0.8 mm thick silicone gaskets (Grace Bio-Labs) onto ultra-clean coverslips. PEG (5 kDa) was conjugated to PLL via a reaction of an amine-reactive succinimidyl valeric acid (SVA) group on PEG and primary amines in PLL. 2% of the 5 kDa PEG had a covalently attached biotin group. Wells were incubated in a 2% biotinylated PLL-PEG solution for 20 min, then washed 8x. Wells were then incubated in a 0.2 mg/ml neutravidin solution for 10 min and washed 8x. GUVs were added to wells at an 8x dilution for 10 min, then washed gently 8x.

### Multibilayers

Lipid multibilayers were prepared according to previously described methods^53^. A lipid mixture of 98 mol% DOPC, 15 mol% DOPS, and 2 mol% DOGS-NTA-Ni was dried to a lipid film under nitrogen gas stream, then dissolved in a 97% hexanes, 3% methanol solution to a concentration of 3.7 mg/ml. 40 μl of this lipid solution was then spin-coated onto a clean glass coverslip at 3000 RPM for 40 s. Spin-coated lipid films were dried under vacuum for 2 hr, then hydrated in silicone gasket wells in the experimental buffer of 20 mM Tris-HCl, 150 mM NaCl, 5 mM TCEP, pH 7.5 at 55°C for 15 minutes, then gradually cooled to room temperature. Wells were washed 6x with buffer before protein was added.

### Cell Culture

Human-derived SUM159 cells gene-edited to add HaloTag (Promega) to both alleles of AP-2 σ2 were a generous gift from Tom Kirchhausen^35^. Cells were grown in 1:1 DMEM high glucose:Ham’s F-12 (Hyclone, GE Healthcare) supplemented with 5% fetal bovine serum (Hyclone), Pen/Strep/L-glutamine (Hyclone), 1 μg/ml hydrocortisone (H4001; Sigma-Aldrich), 5 μg/ml insulin (I6634; Sigma-Aldrich), and 10 mM HEPES (Fisher), pH 7.4 and incubated at 37°C with 5% CO_2_. Cells were seeded onto acid-washed coverslips at a density of 3×10^4^ cells per coverslip for 24 h before transfection with 1-3 μg of plasmid DNA using 3 μL Fugene HD transfection reagent (Promega). HaloTagged AP-2 σ2 was visualized by adding Janelia Fluor 646-HaloTag ligand, a gift from Luke Lavis^36^. 100 nM ligand was added to cells and incubated at 37°C for 15 minutes. Cells were washed with fresh media and imaged immediately.

### Gene Editing

SUM159 AP-2 σ2-HaloTag cells were further gene-edited to knock out both alleles of endogenous Eps15 using clustered regularly interspaced short palindromic repeats (CRISPR)/Cas9. Guide RNA sequences were generated by annealing forward oligo 5’ ^P^CACCG<u>TTGATACAGGCAATACTGGA</u> 3’ and reverse oligo 5’ ^P^AAAC<u>TCCAGTATTGCCTGTATCAA</u>C 3’. The underlined guide sequence targeted bp 77-96 in Eps15. The oligo pair was ligated into pSpCas9(BB)-2A-GFP (Addgene plasmid #48138), a gift from Feng Zhang^54^, at the BbsI site and the resulting plasmid was confirmed by DNA sequencing. SUM159 AP-2 σ2-HaloTag cells were transfected with 2.5 μg of this plasmid containing the guide RNA, Cas9, and an EGFP reporter using Fugene HD transfection reagent. After 24 hr, cells were trypsinized and washed into phosphate-buffered saline (PBS) supplemented with 5% FBS. Cells expressing GFP were sorted using fluorescence-activated cell sorting (FACS) into a 96-well plate at a density of 1 cell per well using a Sony MA900 Multi-Application Cell Sorter. Following clonal expansion, genomic DNA was isolated, then a region surrounding the target sequence was amplified by PCR using oligos 5’ GAAGAGGCACATAACATGTGCAACTATTCCC 3’ and 5’ GGGAATCTTGCCACACACTCAAAGACTTG and sequenced to identify insertions or deletions.

Eps15 knock-out was confirmed by Western blot: SUM159 AP-2 σ2-HaloTag control cells and Eps15 knock-out candidate cells were trypsinized, washed in PBS, and resuspended in ice-cold lysis buffer (20 mM Tris, 150 mM NaCl, 1 mM EDTA, 1 mM EGTA, 1% Triton X-100, 1x Pierce protease inhibitor cocktail tablet, pH 7.5). Total protein concentration was quantified by Bradford assay and equal amounts were loaded on a 4-20% SDS-PAGE gel, then transferred to nitrocellulose membrane. Membranes were blocked in Tris-buffered saline with 0.1% Tween 20 and 5% nonfat dried milk for 1 hr. The primary antibody, Eps15 (D3K8R) Rabbit mAb (# 12460, Cell Signaling Technology), was applied at 1:1,000 in TBS-Tween/5% milk at 4°C overnight.

Membranes were washed 5x in TBS-Tween, then incubated with secondary antibody, Anti-rabbit IgG HRP-linked Antibody (#7074, Cell Signaling Technology) at 1:1,500 in TBS-Tween/1% milk at room temperature for 1 hr. Membranes were washed 5x in TBS-Tween, then imaged using a SuperSignal West Pico kit (Thermo) on a Syngene G:Box Chemi XX6.

### Fluorescence Microscopy

Images of GUVs, multibilayers, and protein droplets were collected on an Axio Observer Z1 (Zeiss) with Yokagawa CSU-X1M spinning disc confocal microscope equipped with a 1.4 NA/100x Plan-Apo oil objective and a cooled EMCCD iXon3 897 camera (Andor Technology). For phase diagram experiments, temperature was monitored by a thermistor placed in the protein solution in a sealed chamber to prevent evaporation. Samples were heated from room temperature at a rate of ∼1°C/minute through an aluminum plate affixed to the top of the chamber.

Live cell images, blue light assays, and droplet FRAP time sequences were collected on a TIRF microscope consisting of an Olympus IX73 microscope body, a Photometrics Evolve Delta EMCCD camera, and Olympus 1.4 NA/100x Plan-Apo oil objective. For cell imaging, the objective was heated to produce a sample temperature of 37°C using an objective heater wrap. The TIRF microscope was equipped with 473 nm, 532 nm, and 640 nm lasers with a 635 nm laser for autofocusing. For droplet imaging the objective was used in widefield mode. All live cell imaging was conducted in TIRF mode at the plasma membrane 16 hr after transfection.

For blue light assays, samples were exposed to 0, 10, or 50 μW 473 nm light as measured out of the objective when in widefield mode. Blue light was applied for 500 ms every 2 s for droplet samples and every 3 s for cell samples. Analysis of movies began after one minute of imaging to allow for blue light to take effect. In particular, cell movies were collected over 11 minutes at 3 s intervals, and the last 10 minutes were analyzed. Clathrin-coated structures were detected and tracked using cmeAnalysis software^37^. Here, the point spread function of the data was used to determine the standard deviation of the Gaussian function. AP-2 σ2 signal was used as the master channel to track clathrin-coated structures. Detected structures were analyzed if they persisted in 3 consecutives frames.

## Supplementary Figures

**Supplementary Figure 1:**
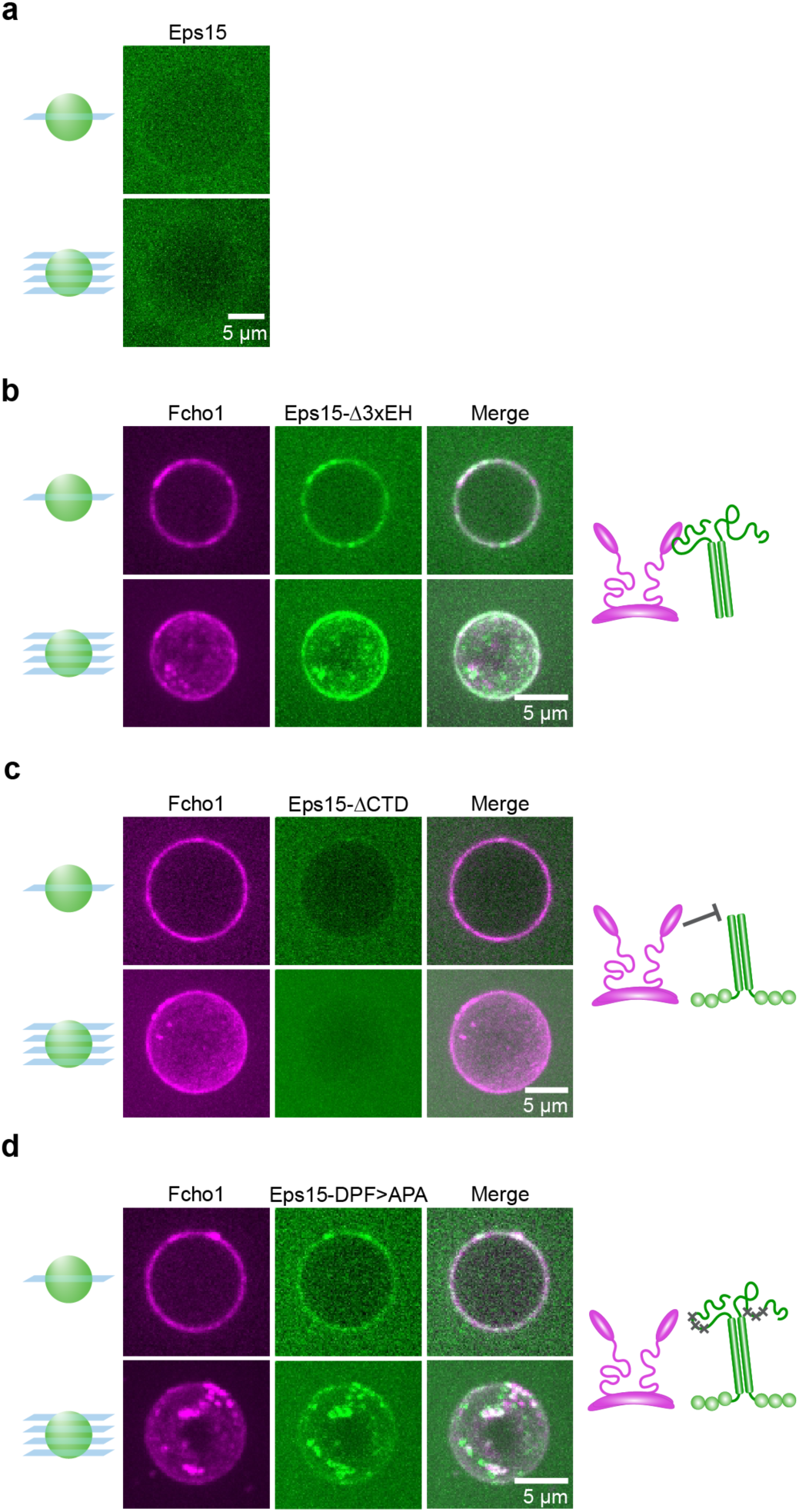
Eps15 mutants and Fcho1 assemble to varying degrees on membrane surfaces. (**a-d**) 500 nM of the indicated proteins were incubated with GUVs containing 79% DOPC, 15% DOPS, 5% PtdIns(4,5)P_2_, and 1% DPEG10-biotin. Upper panels display single center z-slices, lower panels display maximum-projected z-stacks. (**a**) Wild-type Eps15 alone binds weakly and homogeneously to GUVs. (**b-d**) Cartoons depict binding interaction between Fcho1 and Eps15 mutants. (**b**) Fcho1 and Eps15 lacking the EH domains (Eps15-Δ3xEH) occasionally co-assemble into small protein-rich domains. (**c**) Eps15 lacking the C-terminal disordered domain (Eps15-ΔCTD) is not recruited to GUV surfaces by Fcho1. (**d**) Fcho1 and Eps15 containing mutated Fcho1-binding DPF motifs (amino acids 623-636; Eps15-DPF>APA) co-assemble into small protein-rich domains.

**Supplementary Figure 2:**
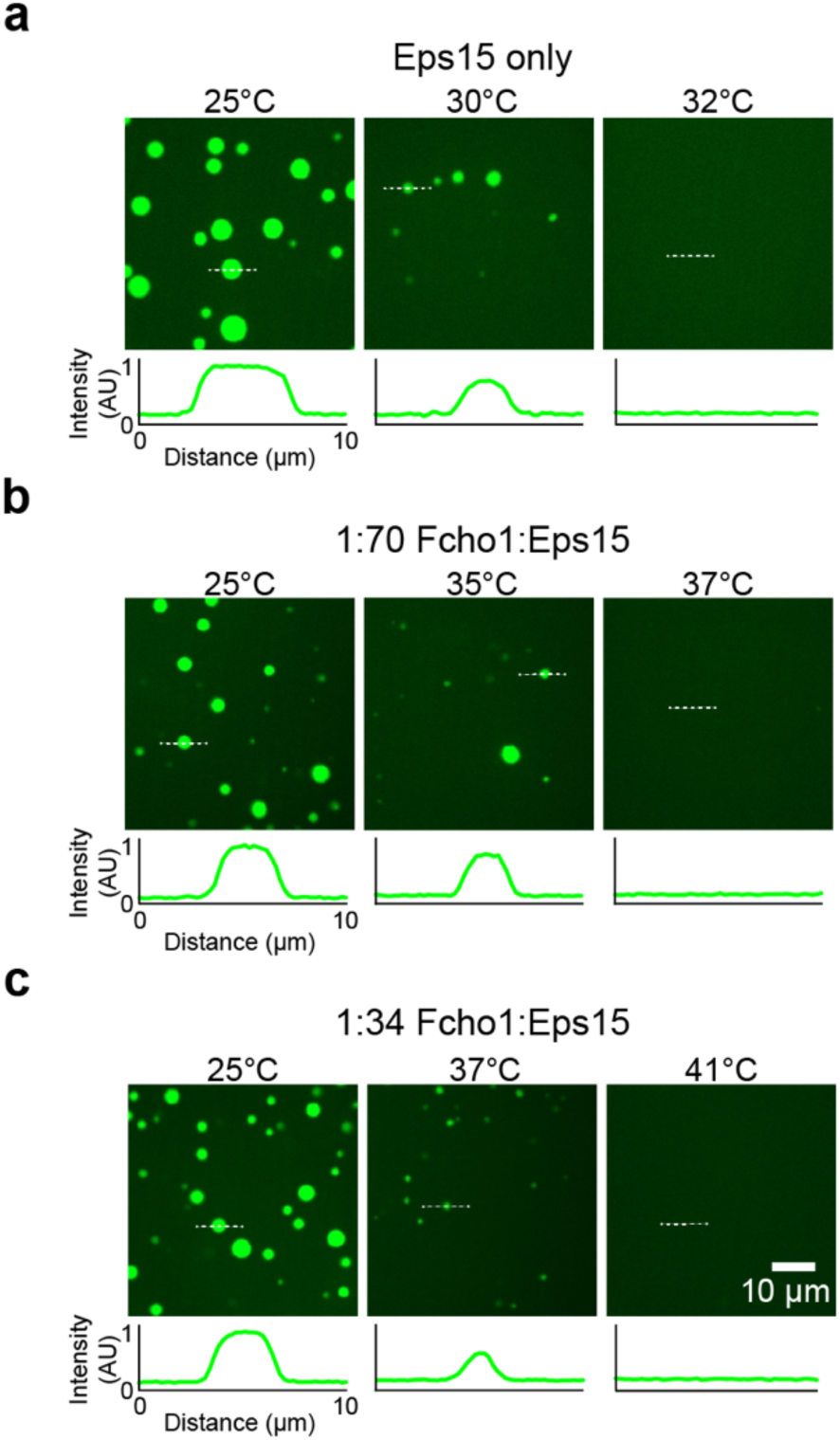
Eps15/Fcho1 droplet:solution partitioning decreases at increasing temperature. (**a-c**) Representative images of protein droplets at increasing temperatures. Plots show fluorescence intensity of Eps15-CF488a measured along dotted lines in each image. Intensity is normalized to the maximum value in the 25°C panel for each set of images. Fcho1 is unlabeled. Total protein concentration is held at 7 μM while Fcho1:Eps15 protein concentration ratio is varied. Droplets are formed from (**a**) 7 μM Eps15, (**b**) 0.1 μM Fcho1, 6.9 μM Eps15, and (**c**) 0.2 μM Fcho1, 6.8 μM Eps15.

**Supplementary Figure 3:**
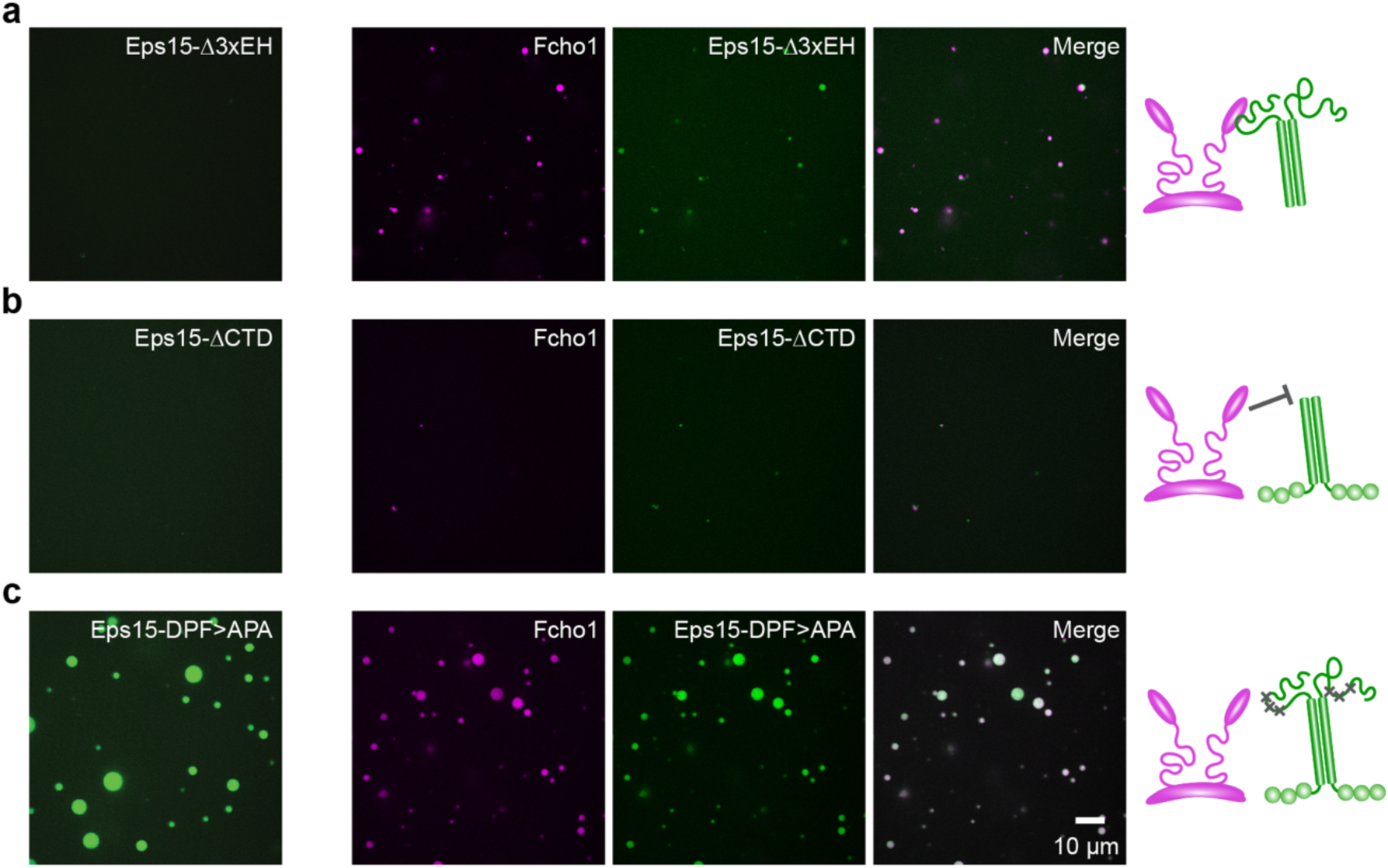
Eps15 mutants and Fcho1 assemble to varying degrees in solution. (**a-c**) Fcho1 is labeled with Atto-594, Eps15 mutants are labeled with CF488a. Proteins were combined at 7 μM total concentration at physiological pH and salt with 3% PEG. Panels on the left show 7 μM Eps15 mutant alone, set of panels on the right show 6.8 μM Eps15 mutant combined with 0.2 μM Fcho1 (34:1). Cartoons depict binding interaction between Fcho1 and Eps15 mutants. (**a**) Eps15 lacking the EH domains (Eps15-Δ3xEH) does not form droplets on its own, but when combined with Fcho1 forms small droplets. (**b**) Eps15 lacking the C-terminal disordered domain (Eps15-ΔCTD) does not form droplets either on its own or when combined with Fcho1. (**c**) Eps15 containing mutated Fcho1-binding DPF motifs (amino acids 623-636; Eps15-DPF>APA) robustly assembles into droplets on its own and co-assembles into droplets with Fcho1.

**Supplementary Figure 4:**
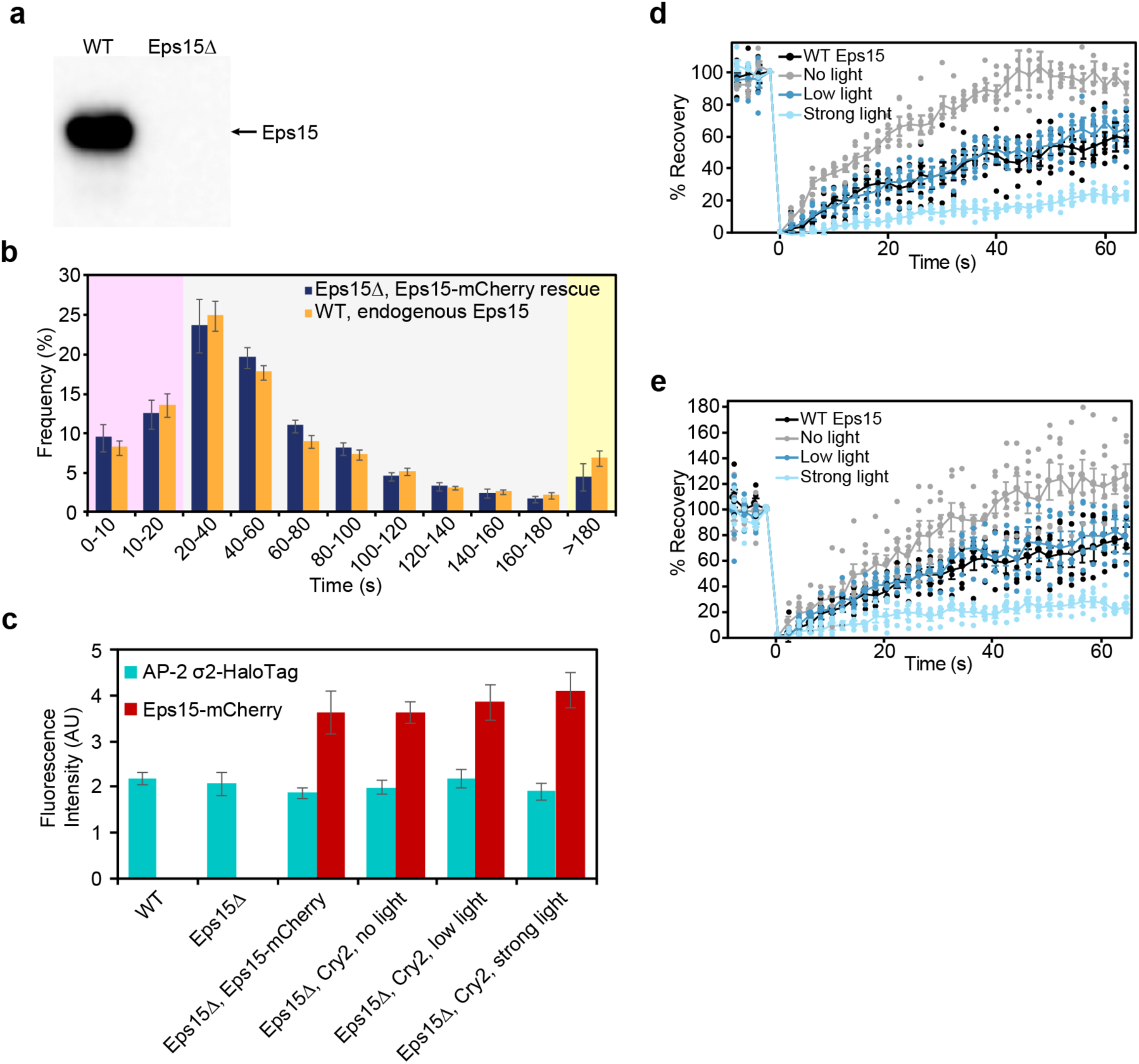
Eps15 knockout cells expressing Eps15-mCherry and in wild-type Eps15 cells are comparable; AP-2 σ2-HaloTag and Eps15-mCherry expression levels are comparable across cell lines in this study. (**a**) Whole cell lysates from WT SUM159/AP-2-HaloTag cells and WT SUM159/AP-2-HaloTag cells gene edited by CRISPR to disrupt Eps15 were separated by SDS-PAGE and immunoblotted for Eps15. (**b**) The lifetime distributions of AP-2 σ2-HaloTag-labeled endocytic structures in Eps15 knockout cells expressing Eps15-mCherry and in wild-type Eps15 cells are nearly identical. (**c**) The average plasma membrane fluorescence intensity of AP-2 σ2-HaloTag::JF_646_ and Eps15-mCherry in the first frame of each movie analyzed in **a** and Figure 4. WT n= 12 cells, Eps15Δ n= 10 cells, Eps15Δ expressing WT Eps15-mCherry n= 11 cells, Eps15Δ expressing Eps15-Cry2 with no blue light n= 14 cells, Eps15Δ expressing Eps15-Cry2 with low blue light n= 19 cells, Eps15Δ expressing Eps15-Cry2 with strong blue light n=13 cells. At least 9,900 endocytic structures were analyzed for each condition. Error bars show SEM. (**d**) Plot from Figure 5d and (**e**) plot from Figure 5f displaying the individual data points that were averaged together for each FRAP curve. n=5-6.

## Supplementary Videos

**Supplementary Video 1: Eps15/Fcho1-rich protein domains merge together on multibilayers.** Video of frames shown in Figure 1h. 100 nM Atto594-labeled Fcho1 (magenta) and 100 nM CF488a-labeled Eps15 (green) were applied to a multibilayer consisting of 73% DOPC, 25% DOPS, and 2% DOGS-NTA-Ni and imaged immediately. Arrowheads indicate merging protein-rich domains. Interval between frames is 3 s, scale bar is 2 μm.

**Supplementary Video 2: Droplets consisting of Eps15 only or Eps15 and Fcho1 merge together while Fcho1 droplets do not merge.** Video of frames shown in Figure 2b, f. Solutions of 7 μM CF488a-labeled Eps15 (upper left), 7 μM Atto594-labeled Fcho1 (lower left), or 6.8 μM CF488a-labeled Eps15 and 0.2 μM dark Fcho1 (right) were prepared. Interval between frames is 0.5 s for Eps15 only and 1 s for Fcho1 only and Eps15/Fcho1, scale bar is 2 μm.

**Supplementary Video 3: Eps15 droplets dissolve uniformly when heated and re**-**form when cooled.** Video on the left shows 7 μM CF488a-labeled Eps15 as it was heated from 30°C to 33°C at a rate of approximately 1°C/min. Video on the right shows 7 μM CF488a-labeled Eps15 as it was allowed to cool from 33°C to 30°C at a rate of approximately 2°C/min. Interval between frames is 1 s, scale bar is 10 μm.

**Supplementary Video 4: Addition of Fcho1 to a single-phase solution of Eps15 induces droplet formation.** Video corresponding to Figure 2h. 3 μM CF488a-labeled Eps15 (green) does not form phase-separated droplets. 0.12 μM Atto594-labeled Fcho1 (magenta) was added at time 0 s and resulted in the formation of droplets that incorporated Eps15 within seconds. Focal plane is near the coverslip, where droplets gradually settled after forming in solution. Video begins 30 s after Fcho1 addition, interval between frames is 1 s, scale bar is 5 μm.

**Supplementary Video 5: Droplets consisting of Eps15-Cry2 and Eps15 show light**-**dependent fusion behavior.** Video of frames shown in Figures 3c and 5a. A solution of 3 μM CF488a-labeled Eps15 (green) and 1 μM Atto594-labeled Eps15-Cry2 (magenta) was exposed to either low blue light (left) or strong blue light (right). Droplets displayed liquid-like merging behavior under low blue light, but failed to merge under strong blue light, indicating solid-like assembly. Interval between frames is 2 s, scale bar is 5 μm.

**Supplementary Video 6: Blue light exposure impacts the dynamics of clathrin**-**coated structures in cells expressing Eps15-Cry2.** Representative movies of cells used for analysis in Figures 4 and 5. Cells were gene-edited to delete endogenous Eps15 and to express AP-2 σ2-HaloTag. AP-2 is labeled with JF_646_ dye and displayed in cyan. Cells express either no Eps15 (“Eps15Δ”), Eps15-mCherry (“WT Eps15”) or Eps15-Cry2-mCherry (“Light off”, “Low light”, and “Strong light”), displayed in red. Movies begin after 1 min of imaging with or without blue light. Interval between frames is 3 s, scale bar is 10 μm.

## Acknowledgements

We thank T. Kirchhausen for the gift of SUM159/AP2σ2-HaloTag cells and L. Lavis for the gift of JF_646_ HaloTag ligand. This research was supported by the National Institutes of Health through grants R01GM120549 to J.C.S. and E.M.L. and F32GM133138 to K.J.D. and by a National Science Foundation Graduate Research Fellowship (DGE-1610403) to G.K. The University of Texas Health Science Center at San Antonio (UTHSCSA) Center for Macromolecular Interactions is supported by the Cancer Therapy and Research Center through the National Cancer Institute P30 Grant CA054174, and Texas State funds provided through the UTHSCSA Office of the Vice President for Research.

## Author Contributions

K.J.D. and J.C.S. designed experiments and wrote and edited the manuscript. K.J.D. performed, and analyzed experiments. G.K., J.B.R., and L.W. expressed and purified proteins. C.C.H. designed and built key equipment and assisted with microscopy. E.M.L. consulted on protein purification. All authors consulted on manuscript preparation and editing.

## Data Availability Statement

All data supporting this work are available on request to the corresponding author.

